# Tumor-specific changes in Kaposi sarcoma-associated herpesvirus genomes in Ugandan adults with Kaposi sarcoma

**DOI:** 10.1101/2020.05.04.076638

**Authors:** Jan Clement Santiago, Jason Goldman, Hong Zhao, Alec Pankow, Fred Okuku, Michael Schmitt, Lennie Chen, C. Alexander Hill, Corey Casper, Warren T. Phipps, James I. Mullins

**Affiliations:** Departments of Microbiology, University of Washington, Seattle, WA US; Departments of Medicine, University of Washington, Seattle, WA US; Fred Hutchinson Cancer Research Center, Seattle, WA US; Uganda Cancer Institute, Kampala, UG; Departments of Global Health, University of Washington, Seattle, WA US; Departments of Laboratory Medicine, University of Washington, Seattle, WA US

## Abstract

Intra-host evolved tumor virus variants have provided insights into the risk, pathogenesis and treatment responses of associated cancers. However, the intra-host variability of Kaposi sarcoma-associated herpesvirus (KSHV), the etiologic agent of Kaposi sarcoma (KS), has not been explored at the whole viral genome level. An accurate and detailed description of KSHV intra-host diversity in whole KSHV genomes from matching tumors and oral swabs from Ugandan adults with HIV-associated KS was obtained by deep, short read sequencing, using duplex unique molecular identifiers (dUMI) – random double-stranded oligonucleotides that barcode individual DNA molecules before library amplification. This allowed suppression of PCR and sequencing errors down to ∼10^−9^/base. KSHV genomes were assembled *de novo*, and identified rearrangements were confirmed by PCR. 131-kb KSHV genome sequences, excluding major repeat regions and averaging 2.3 × 10^4^ reads/base, were successfully obtained from 23 specimens from 9 individuals, including 7 tumor-oral pairs. Sampling more than 100 viral genomes in at least one specimen per individual showed that KSHV genomes were virtually homogeneous within samples and within individuals at the point mutational level. Heterogeneity, if present, was due to point mutations and genomic rearrangements in tumors. In 2 individuals, the same mutations were found in distinct KS tumors. The K8.1 gene was inactivated in tumors from 3 individuals, and all KSHV genomic aberrations retained the region surrounding the first major internal repeat (IR1). These findings suggest that lytic gene alterations may contribute to KS tumorigenesis or persistence.

**Author summary:** Kaposi sarcoma (KS) is a leading cancer in sub-Saharan Africa and in those with HIV co-infection. Infection by Kaposi sarcoma-associated herpesvirus (KSHV) is necessary for KS, yet why only few KSHV infections develop into KS is largely unknown. While strain differences or mutations in other tumor viruses are known to affect the risk and progression of their associated cancers, whether KSHV genetic variation is important to the natural history of KS is unclear. Most studies of KSHV diversity have characterized only ∼4% of its 165-kb genome and may have been impacted by PCR or cloning artifacts. Here, we performed highly sensitive, single-molecule sequencing of whole KSHV genomes in paired KS tumors and oral swabs from 9 individuals with KS. We found that KSHV genomes were virtually identical within individuals, with no evidence of quasispecies formation nor multistrain infection. However, KSHV genome aberrations and inactivating mutations appeared to be a common, tumor-associated phenomenon, with some mutations shared by distinct tumors within an individual. Certain regions of the KSHV genome featured prominently among tumor-associated mutations, suggesting that they are important contributors to the pathogenesis or persistence of KS.

## Introduction

Kaposi Sarcoma (KS) is one of the most common cancers of HIV-infected individuals [1,2], with the burden of disease disproportionately borne by people in sub-Saharan Africa [3]. A gamma herpesvirus, Kaposi sarcoma-associated herpesvirus (KSHV), is the etiologic agent of KS and consistently detected in tumor tissues [4,5]. KSHV also can be shed in saliva, thought to be a primary mode of transmission [6–8]. Only a small fraction of KSHV infections progress to KS, and the factors contributing to KS pathogenesis are poorly understood. The development of KS is associated with HIV infection and immunosuppression [9], but others factors, including KSHV genome variation, may contribute to differential outcomes of KSHV infection.

Studies of other human oncogenic viruses reveal that viral genetic variation or *de novo* mutations may be important to their pathogenicity, as is the case for cancers associated with human papilloma viruses and Merkel cell polyomavirus [10]. Epstein Bar virus (EBV), another gamma-herpesvirus like KSHV, is associated with a variety of neoplasms. In EBV infections, intra-host evolved viruses may have a role in pathogenesis of associated cancers [11–13]. Additionally, EBV strains isolated from nasopharyngeal carcinoma (NPC) biopsies have unique genomic [14,15] and phenotypic [16–18] variations compared to other strains and isolates from geographically clustered individuals without cancer [19]. NPC-associated strains were also found to have increased tropism for epithelial-versus B-cells [16,20]. These findings suggest that viral genetic heterogeneity can affect EBV virulence. Whether KSHV genetic variation can similarly influence KS pathogenesis or manifestation is unknown.

KSHV encodes oncogenes that dysregulate cell cycle, cell-to-cell adhesion, inflammation and angiogenesis [21–24], and heterogeneity in these genes might result in differences in KSHV infection outcomes and clinical manifestations. For example, polymorphisms in the microRNA region of KSHV have been correlated with the development of multicentric Castleman disease and KSHV-associated inflammatory cytokine syndrome with and without KS [25]. The K1 gene, the most variable KSHV gene, is conventionally used for KSHV strain subtyping. Subtype A has been associated with more aggressive KS than subtype C [26,27], while subtype B has been associated with better KS prognoses [28]. However, correlations of KSHV genetic subtypes with virulence has not been consistently observed [29–32].

KSHV whole genome sequences provide a far more comprehensive picture of KSHV diversity than the KSHV variable regions alone. Most of the publicly available KSHV genomes have been reported in only the last 5 years [33–37], and they demonstrated that polymorphisms in the ∼130-kb KSHV non-repeat genomic region outside of the K1 gene contribute much more to KSHV diversity than the 0.9 kb hyper-variable K1 gene by itself [33,34]. KSHV genomes are, moreover, replete with signatures of recombination [36], potentially complicating disease risk association solely with K1 subtypes. Principal component analysis of 70 KSHV genomes, all from different individuals, failed to reveal strain specificity for the development of KS [36].

Whether KSHV results in significant intra-host diversity is unclear. Some studies examining KSHV in different compartments or multiple clones from a single individual have reported KSHV quasispecies, multi-strain infections [38–42] and intra-host evolution [37,40,43], while only a single persisting strain is typically found within individuals with AIDS-associated KS [44–47]. The existence of KSHV intertypic recombinants [36,44,48–50] indicate that co-infection of divergent KSHV strains must be occurring at least sporadically.

The assessment of intra-host diversity can be easily biased by artifacts introduced during sample preparation and when sequencing from PCR products [45]. Short read, next generation sequencing can have high error rates due to PCR misincorporation, end-repair artifacts, insufficient sequencing depth, and DNA damage from long, repeated high-temperature incubations during PCR and enrichment reactions [51–54]. These methodological limitations can undermine interpretation of observed intra-host KSHV variations.

To accurately assess KSHV intra-host polymorphisms associated with KS tumors and detect minor KSHV sequence variants, we sequenced KSHV genomes using a highly sensitive short-read sequencing method termed “duplex sequencing” [55]. This method incorporates duplex unique molecular identifiers (dUMI), which are double-stranded strings of random base pairs to barcode individual DNA molecules before PCR amplification and RNA-bait enrichment [55]. By utilizing dUMI-consensus reads of each DNA molecule in a sample library, PCR-associated errors are reduced to ∼10^−9^, revealing the original sequence variation within a sample library [55]. In the present study, we report the results of duplex sequencing of whole KSHV genomes in paired tumors and oral swabs of 9 Ugandan adults with HIV-associated KS.

## Methods

### Study Cohort and Specimen collection

Specimens were obtained from participants enrolled in the “HIPPOS” Study, an ongoing prospective cohort study, begun in 2012, of KS patients initiating treatment at the Uganda Cancer Institute (UCI) in Kampala, Uganda. This protocol was approved by the Fred Hutchinson Cancer Research Center Institutional Review Board, the Makerere University School of Medicine Research and Ethics Committee (SOMREC), and the Uganda National Council on Science and Technology (UNCST). All participants provided written informed consent.

Participants were eligible for the HIPPOS study if they were >18 years of age with biopsy-proven KS, and ART- and chemotherapy-naïve at enrollment. Participants attended 12 study visits over a one-year period and received treatment for KS consisting of ART and chemotherapy (combination bleomycin and vincristine or paclitaxel) per standard protocols by UCI physicians. At each visit, participants received a detailed physical exam to assess clinical response using the ACTG KS response criteria [56].

Participants provided plasma samples at each visit for KSHV, CD4 and HIV viral load testing, and in addition, provided up to 9 biopsies of KS lesions before, during, and after KS treatment. KS tumor biopsies were obtained using 4mm punch biopsy tools after cleaning the skin with alcohol, and either snap-frozen at the clinic site and stored in liquid nitrogen (LN2) or placed in RNAlater and stored at −80^0^C. Study clinicians collected swabs of the oral mucosa at each study visit and participants self-collected oral swabs at home for 1 week after the visit after education on the sample collection technique, as has been previously validated by our group in Uganda [57]. Briefly, a Dacron swab is inserted into the mouth and vigorously rubbed along the buccal mucosa, gums, and hard palate. The swab is then placed in 1 mL of filter-sterilized digestion buffer [58] and stored at ambient temperature [59] before being placed at −20°C for storage.

### DNA preparation

DNA was extracted from 300µL homogenized tumor lysates using the AllPrep DNA/RNA Mini Kit (QIAGEN, Cat. # 80204) and eluted into 100µL EB Buffer. For oral swab specimens, DNA was extracted from the swab tip eluate using the QIAamp Mini Kit (Qiagen, Cat.# 51304) following the manufacturer’s protocol. Purification of DNA from saliva stabilized in RNAprotect® Saliva Reagent (Qiagen) was performed following the manufacturer’s protocol with the following modifications: there was no initial pelleting or PBS wash, 20 µL proteinase K was used per 200 µL specimen, and DNA was eluted in 50 µL water. DNA was quantified using a NanoDrop™ 1000 Spectrophotometer (ThermoScientific).

### PCR

All PCR preparations were done in a PCR-clean room, except for the addition of control templates. PCR was conducted using the PrimeSTAR GXL kit (Takara, Cat. # R050B) with ThermaStop™ (Thermagenix) added. Cycling conditions were: 98°C for 2 mins; 35 cycles of 98°C for 10 secs, 60-65°C (depending on primer) for 15 secs, 68°C for 1min/kb; 68°C for 3 mins and then hold at 4°C. Primer sequences are listed in **S1 Table**.

### Copy number quantification

KSHV genome copy numbers were quantified by digital droplet PCR (ddPCR) using the QX200™ Droplet Generator and Reader (Bio-Rad), with ddPCR™ SuperMix for Probes No dUTP (Bio-Rad, Cat. # 186-3024). Primers and probes were designed (**S1 Table**) to detect 4 KSHV-unique genes K2/vIL-6, ORF16/vBCL-2, ORF50/RTA and ORF73/LANA (**Fig 1A**), and the KSHV genome copy numbers reported were the average of the 4 measures. 420 ng BCBL-1 cell line DNA diluted 1:10,000, about 475 genome copies, was used as positive control. 1 ng human genomic DNA (Bioline, Cat. # BIO-35025) was used as negative control, and water as a no template control. Cycling conditions were: 95°C for 10 mins; 40 cycles of: 94°C for 30 secs, 56°C for 30 secs, 60°C for 1 min; one cycle at 98°C 10 for mins, and then hold at 12°C. The KSHV on-target percent was calculated using the copy number quantification by ddPCR normalized to the total nucleic acid concentration.

**Figure 1.**
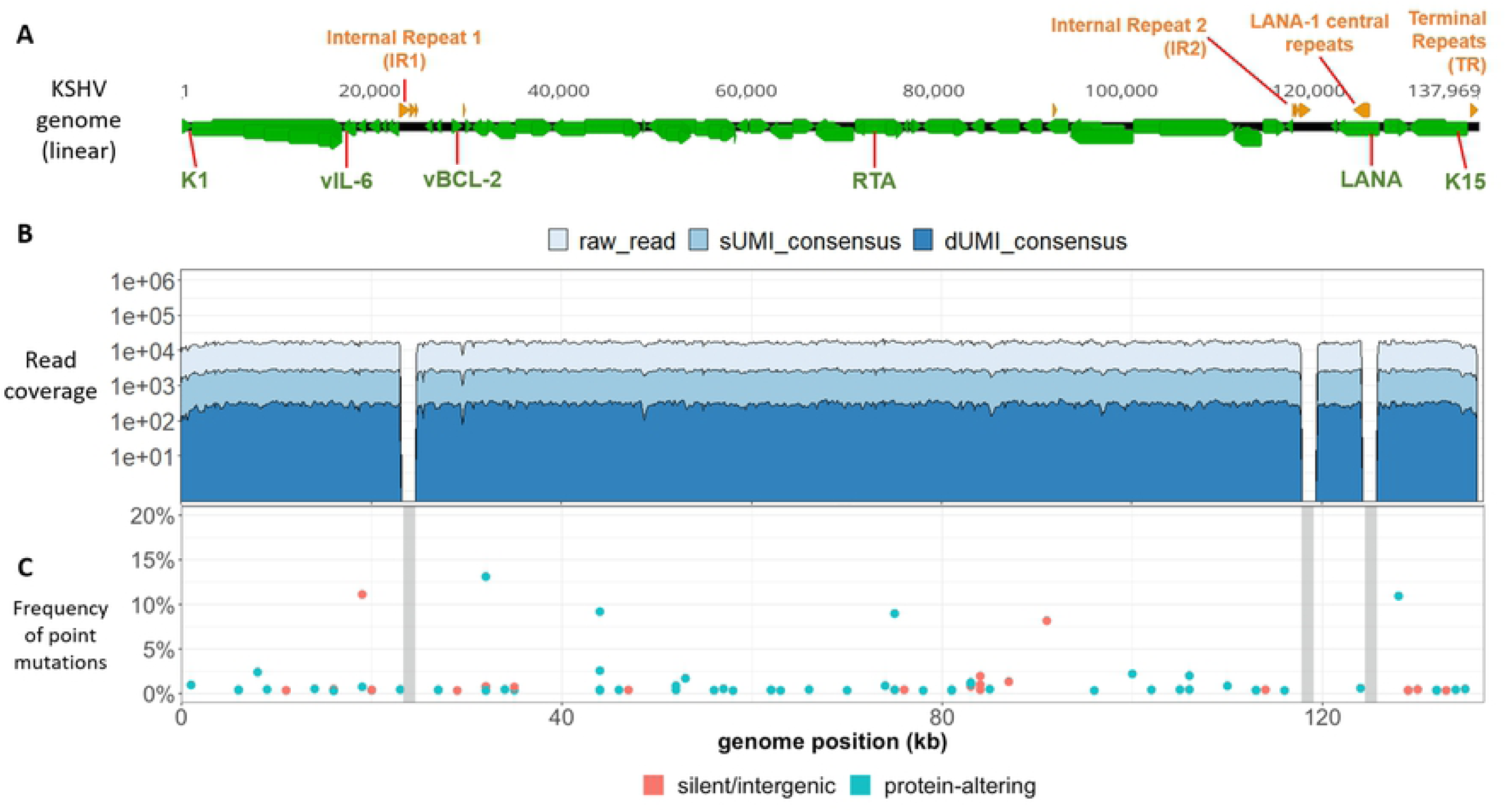
KSHV genomes in BCBL-1 cells have low point mutational diversity. (A) Schematic representation of a linear KSHV genome, with genes colored in green and the major repeat regions in orange. The locations of the K1, vIL-6, vBCL-2, RTA, LANA and K15 genes used for genome quantitation are indicated. (B) Raw (light blue), sUMI-consensus (blue) and dUMI-consensus (dark blue) read coverage along the *de novo* assembled, BCBL-1 KSHV genome. Major repeat regions were masked (gray columns). (C) Bubble plot of minor sequence variants. Each bubble represents a position within the genome at which a variant base or indel was detected, colored by whether they were predicted to be silent or protein-altering mutations. Mutations likely to be silent included synonymous and intergenic point mutations, while protein-altering mutations included non-synonymous, nonsense and frameshift mutations. Bubble height represents variant base frequency among dUMI-consensus reads at that position. Vertical grey columns represent the masked repeat regions.

### UMI-addition and library preparation (Fig 2)

To obtain ∼500-bp DNA fragments, 10-20 ng/μL of DNA extract in 100 µL chilled TLE buffer (10mM Tris, pH8.0, 0.1mM EDTA) was sheared using a Bioruptor™ (Diagenode) on high power for up to 15 min. Fragment sizes were assessed on 1.5% agarose gels. Sheared DNA was bead-purified using 1.2X volume of Agencourt AMPure XP Beads (Beckman Coulter Cat. # A63880) and eluted in 50 µL water. Library preparation (end repair, A-tailing and adapter-ligation) was performed using the KAPA HyperPrep Library Preparation Kit (Cat. # KR0961/KK8503). Double-stranded DNA adapters contained a random 12-bp dUMI sequence and a defined 5-bp spacer sequence added to Illumina TruSeq adaptor sequences [60] (**S1 Fig**). Subsequently, DNA was bead-purified with 1X volume of beads and eluted in 50 µL water.

**Figure 2.**
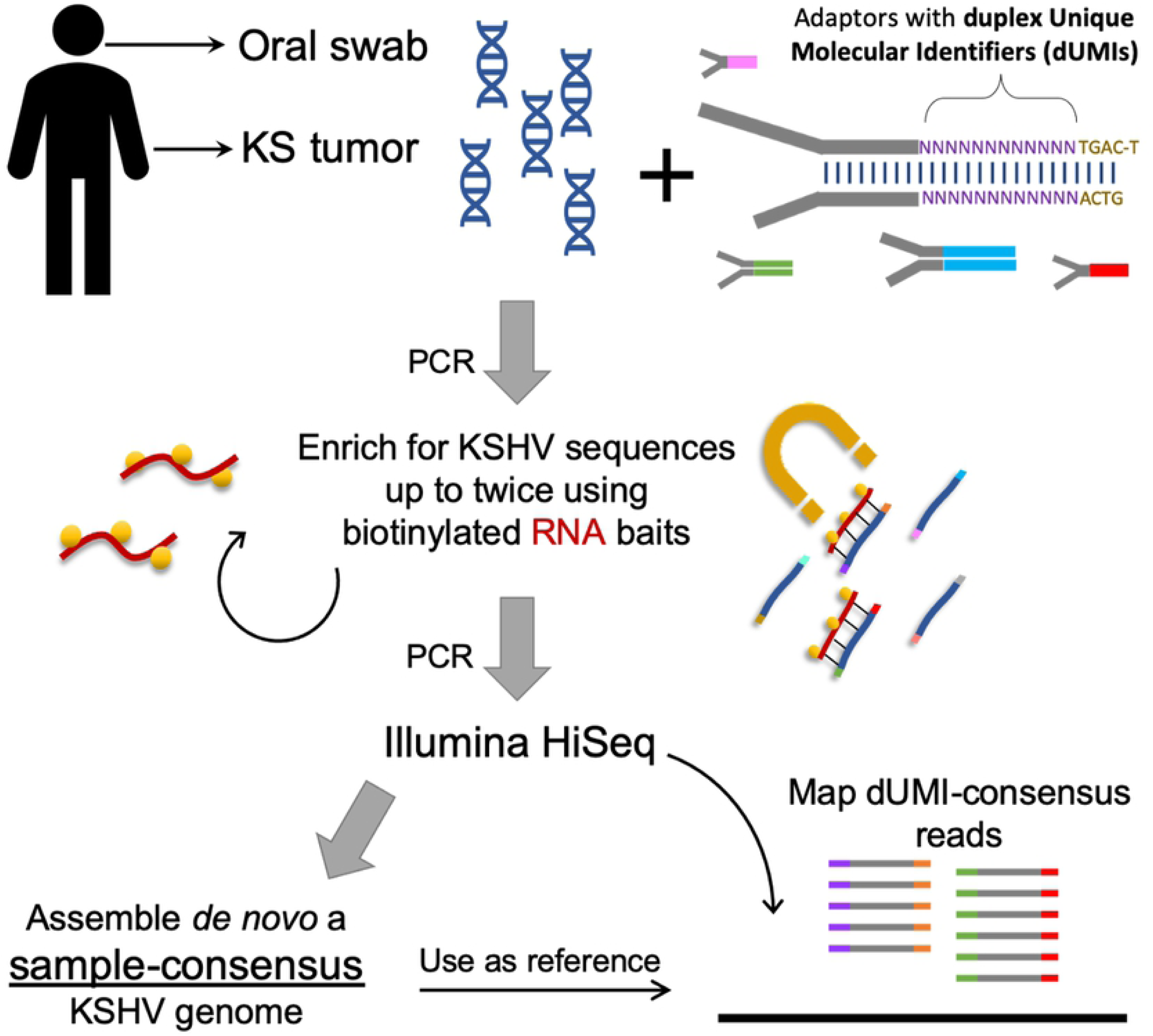
Workflow for analyzing intra-host KSHV genome diversity from clinical samples. Each study participant contributed KS tumors and oral swabs. Sequencing libraries were prepared from DNA extracts from each sample with adaptors containing duplex Unique Molecular Identifiers (dUMIs). Adaptor-labelled DNA libraries were enriched using biotinylated RNA baits homologous to KSHV sequences. Captured DNA was PCR-amplified to levels sufficient for Illumina HiSeq sequencing. For most samples, libraries were subjected to a second round of enrichment and PCR amplification. Upon sequencing, whole KSHV genomes were first assembled *de nov*o from each sample without the use of dUMIs. The sample-specific genomes generated (sample-consensus) were then used as reference to map the consensus of reads with identical dUMI-tags (i.e., dUMI-consensus reads).

DNA libraries were subjected to pre-enrichment amplification with primers mws13 & mws20 (**S1 Table, S1 Fig**) and KAPA HiFi Hot Start polymerase. PCR conditions were: 95°C 4 mins; 5-8 cycles of 98°C 20 sec, 60°C 45 sec, 72°C 45 sec; 72°C 3 mins, 4°C hold. If the bead-purified elution from the end repair and adapter step had more than 240 ng total, it was divided into 50 µL PCR reactions of ≤240 ng and pooled after amplification. PCR products were then bead-purified as above with 1.2X volume beads and elution in 100 µL water, quantified with Nanodrop, and their sizes assessed using a Bio-analyzer (Agilent DNA 7500) or Qiaxcel (QIAGEN AM420).

### Library enrichment & sequencing

Biotinylated RNA baits for enriching KSHV sequences in the library were those designed in [61] and were obtained from Agilent, Inc. (Santa Clara, CA). The design was a 120-bp, 12X tiling of the genome of KSHV isolate GK18 (Genbank ID: AF148805.2). The diversity of the bait library was further increased by including K1, ORF75, K15, ORF26 and TR sequences of strains JSC-1 (Genbank ID: GQ994935.1), DG1 (Genbank ID: JQ619843.1), BC-1 (Genbank ID: U75698.1), BCBL-1 (Genbank ID: HQ404500.1), Sau3A (Genbank ID: U93872.2), and all Western and African strain sequences in [29,33] (Genbank ID: AF130259, AF130266, AF130267, AF130281, AF130305, AF133039, AF133040, AF133043, AF133044, AF151687, AF171057, AF178780, AF178810, AF220292, AF220293, AY329032, KT271453, KT271454, KT271455, KT271456, KT271457, KT271458, KT271459, KT271460, KT271461, KT271462, KT271463, KT271464, KT271465, KT271466, KT271467, KT271468).

Target enrichment was performed using SureSelect Target Enrichment Kit v1.7 (Agilent) with all suggested volumes reduced by half. DNA hybridized to biotinylated-RNA baits was captured with streptavidin beads (Dynabeads MyOne Streptavidin T1, Invitrogen) and resuspended in 20µL water. The DNA-streptavidin bead mixture was used directly in post-enrichment PCR amplification with primers mws13 and mws21, the latter of which includes a sample index sequence (**S1 Table, S1 Fig**). The PCR cycle number ranged from 10-16, with products monitored every 2 to 3 cycles on a TapeStation (Agilent) to ensure correct fragment sizes (∼500bp). When over-amplification resulted in library fragment sizes much larger than expected, a single “reconditioning” PCR cycle with fresh reagents was done [62]. PCR products were cleaned using 1.2X volume AMPure XP beads and the eluted DNA library was sequenced using Illumina HiSeq 2500 with 100-bp paired end reads. For some tumor samples with low KSHV copy numbers and all oral swab samples, a second library enrichment was performed.

### *De novo* assembly of sample-consensus genomes

Initially, a sample-consensus KSHV genome (**Fig 2**) was generated *de novo* from paired-end reads of each sample using custom scripts (**S2 Fig**, https://github.com/MullinsLab/HHV8-assembly-SPAdes). At this stage, the first 17-bp from read ends were trimmed to remove dUMI sequences. Next, reads were subjected to windowed quality filtering using *sickle pe* [63] with a quality threshold of 30 and a window size 10% of read length. Filtered reads were aligned to a human genome (GRCh38 p12, GenBank GCA_000001405.27) using *bwa mem* [64]. Unmapped reads were used as input into the de novo assembler SPAdes v3.11.1 [65], with the setting -*k 21,35,55,71,81*. This oftentimes yielded 3 to 4 scaffolds that together encompassed the entire 131-kb unique sequence regions of KSHV, bounded by the major repeat regions: Internal Repeat 1 (IR1), Internal Repeat 2 (IR2), LANA central repeat and Terminal Repeats (TR) (**Fig 1A**). Next, all scaffolds over 500 bp were aligned using *bwa mem* to the genome of reference KSHV strain GK18. From the aligned scaffolds a draft genome was generated in Geneious (Biomatters, Ltd) with manual correction as needed. To finish the assembly, GapFiller v1.1 [66] was used, setting *bwa* as the aligner and filtered paired-end reads as the input library. The genomes were annotated in Geneious from the GK18 reference, also adding the T1.4 annotation based on [67]. The major repeat regions were masked with Ns since they were poorly resolved by assembly of short reads that can map to multiple locations within the repeat regions.

### Variant identification from dUMI-consensus reads

Paired-end reads, including their dUMI sequence tags, were mapped by *bwa* to sample-consensus genomes (**Fig 2**) using a Makefile adapted from [60] (https://github.com/MullinsLab/Duplex-Sequencing). Briefly, all reads mapping to the same genomic position were collapsed by single strand UMIs (sUMI) to make sUMI-consensus reads (**S2 Fig**). Complementary UMI tags from opposing strands were matched to create dUMI-consensus reads, removing nearly all PCR polymerase misincorporation and chimera artifacts. Nine bases from both read ends were then trimmed to minimize read end artifacts. Discrepancies between mapped dUMI-consensus reads and the sample-consensus genomes were manually inspected and corrected in Geneious as needed. Only the remaining discrepancies were considered to be sequence variants that existed prior to PCR amplification.

All genome and subgenome sequence alignments were done using MAFFT [68] [algorithm FFT-NS-i x1000, scoring matrix 1PAM/k=2], and all phylogenetic trees were inferred using RAxML [69] (-f d, GTR gamma, N=100 starting trees), using a representative KSHV genome from each individual. The NeighborNet phylogenetic network was generated using SplitsTree5, excluding gap sites [70].

Consensus genome sequences were deposited in GenBank (Accession numbers: XXX) with coordinates of rearrangements, when present, indicated.

### Integration analysis

Systematic searches for KSHV integration into the human genome were done in two ways. First, each library was searched using local BLASTN against both human and KSHV sequences and then using the Perl script SummonChimera [71] to extract coordinates of potential integration sites. Second, a sample-consensus KSHV genome was appended as an extra chromosome to the human genome reference GRCh38 p12. The appended human genome reference was used to map sUMI consensus reads via Speedseq [72] to generate alignment files with only discordant or split reads. These were input into LUMPY for structural variant analysis [73]. Human chromosomes linked to KSHV sequences were taken to be putative integration sites.

## Results

### Assessment of the dUMI sequencing protocol with a KSHV infected cell line

As part of the optimization of the dUMI-sequencing protocol, KSHV genome sequences were first obtained from an early passage of BCBL-1, a KSHV-infected PEL cell line [74]. BCBL-1 cells were grown as previously described [75]. After DNA extraction, KSHV sequences corresponded to ∼0.16% of the total DNA using a ddPCR assay for ORF73 and T0.7-K12, and normalized by comparison to the human gene POLG. Following a single round of bait capture, the fraction of sequence reads corresponding to KSHV from BCBL-1 DNA extracts (i.e., the “on-target” level), was 15.6%, corresponding to 173-fold enrichment.

Sequencing of the BCBL-1 KSHV genome produced a mean coverage of >10,000 reads per base excluding the repeat regions. Collapsing raw reads by identical sUMI to generate sUMI-consensus reads results in a mean of 2,552 sUMI reads per base. When collapsed further into consensus sequences derived from both strands, a mean of 286 dUMI reads per base was obtained that was essentially free of PCR errors (**Table 1; Fig 2B**). Since each dUMI tags a unique DNA molecule before PCR, the number of unique dUMI tags indicates the number of unique viral templates sequenced [55,76]. Using this measure, 286 also approximates the number of KSHV genomes sampled from BCBL-1.

**Table 1.** Origin and processing results from specimens for KSHV genome analysis.

| Pt ID | Age | Sex | Plasma HIV RNA (copies/mL) | CD4 T cells/μL | Sample ID | Sample Type | % On-target | | Times enriched | Fold enrichment | Mean read coverage | Mean dUMI-consensus read coverage | Genome length (excluding repeats and Ns) | # v |
| --- | --- | --- | --- | --- | --- | --- | --- | --- | --- | --- | --- | --- | --- | --- |
|  |  |  |  |  |  |  | Pre-enrichment | Post-enrichment |  |  |  |  |  |  |
| n/a | n/a | n/a | n/a | n/a | BCBL-1 | cell line | 0.09% | 15.61% | 1x | 173 | 7,586 | 286 | 132,676 |  |
| U003 | 25 | M | 759,635 | 45 | **U003-C** | **Tumor** | **0.37%** | **7.15%** | **1x** | **1,925** | **15,635** | **259** | **131,292** |  |
|  |  |  |  |  | *U003-o1* | *Oral swab* | <0.01% | 35.20% | 2x | 2,919,559 | 47,445 | 7 | 131,102 |  |
|  |  |  |  |  | *U003-o2* | *Oral swab* | 0.03% | 31.60% | 2x | 120,023 | 63,258 | 292 | 131,129 |  |
|  |  |  |  |  | *U003-o3* | *Oral swab* | <0.01% | 41.90% | 2x | 1,007,251 | 58,467 | 41 | 131,143 |  |
| U004 | 37 | M | 277,655 | 85 | **U004-C** | **Tumor** | **0.16%** | **17.20%** | **1x** | **10,567** | **18,787** | **32** | **131,277** |  |
|  |  |  |  |  | **U004-D** | **Tumor** | **0.17%** | **75.10%** | **1x** | **45,228** | **58,120** | **1,263** | **131,237** |  |
|  |  |  |  |  | *U004-o1* | *Oral swab* | 0.01% | 18.90% | 2x | 335,500 | 36,466 | 55 | 131,277 |  |
| U007 | 26 | M | 91,096 | 136 | **U007-B** | **Tumor** | **0.06%** | **86.90%** | **2x** | **137,642** | **67,373** | **1,040** | **131,352** |  |
|  |  |  |  |  | *U007-o1* | *Oral swab* | 0.01% | 27.60% | 2x | 402,068 | 41,922 | 198 | 131,126 |  |
| U008 | 56 | M | 860,937 | 488 | **U008-B** | **Tumor** | **0.49%** | **16.97%** | **1x** | **3,470** | **19,553** | **644** | **131,142** |  |
|  |  |  |  |  | **U008-D** | **Tumor** | **0.40%** | **29.98%** | **2x** | **7,534** | **25,341** | **188** | **131,102** |  |
|  |  |  |  |  | *U008-o1* | *Oral swab* | <0.01% | 23.50% | 2x | 846,462 | 46,611 | 111 | 131,116 |  |
| U020 | 27 | M | 118,191 | 370 | **U020-B** | **Tumor** | **0.70%** | **1.47%** | **1x** | **210** | **12,774** | **154** | **131,102** |  |
|  |  |  |  |  | **U020-C** | **Tumor** | **0.03%** | **34.38%** | **2x** | **98,720** | **15,438** | **0** | **131,471** |  |
|  |  |  |  |  | *U020-o1* | *Oral swab* | <0.01% | 77.30% | 2x | 3,090,161 | 7,807 | 80 | 131,115 |  |
|  |  |  |  |  | *U020-o2* | *Oral swab* | <0.01% | 51.40% | 2x | 3,702,047 | 6 | 0 | 128,518 |  |
| U023 | 33 | F | 338,285 | 191 | *U023-o1* | Oral | 0.01% | 2.70% | 2x | 25,189 | 19,877 | 1 | 131,122 |  |
| U030 | 40 | M | 100,184 | 70 | **U030-C** | **Tumor** | **1.35%** | **15.68%** | **1x** | **1,162** | **36,741** | **468** | **131,282** |  |
|  |  |  |  |  | *U030-o1* | *Oral swab* | <0.01% | 5.40% | 2x | 196 | 9,402 | 0 | 128,676 |  |
| U032 | 23 | F | 587,149 | 274 | **U032-B** | **Tumor** | **0.03%** | **43.50%** | **2x** | **127,887** | **66,595** | **854** | **131,266** |  |
|  |  |  |  |  | *U032-o1* | *Oral* | <0.01% | 7.70% | 2x | 4,422,546 | 7,303 | 1 | 130,842 |  |
| U034 | 47 | F | 130,375 | 237 | **U034-B** | **Tumor** | **0.14%** | **84.40%** | **2x** | **41,282** | **71,473** | **1,653** | **131,248** |  |
|  |  |  |  |  | **U034-C** | **Tumor** | **0.13%** | **30.70%** | **2x** | **22,056** | **17,074** | **126** | **131,088** |  |
|  |  |  |  |  | *U034-o1* | *Oral swab* | <0.01% | 7.30% | 2x | 3,073,915 | 7,635 | 2 | 130,754 |  |
|  |  |  |  |  | *U034-o2* | *Oral swab* | <0.01% | 6.00% | 2x | 6,411,538 | 4,087 | 1 | 130,884 |  |

Eighty-one base positions (0.06%) in the BCBL1 consensus KSHV genome had detectable variants in dUMI-consensus reads, and the average frequency of minor variants was 1.35%. No variant exceeded 14% of the total dUMI-consensus reads at any position (**Fig 1C)**. No doubling of read coverage was found within the 19-kb genomic region previously reported in the BCBL-1-derived KSHV recombinant clone BAC-36 [77].

The consensus, *de novo*-assembled KSHV genome in BCBL-1 had 3 differences from the published BAC-36 sequence: a C→A change in the noncoding sequence before ORF K5 (BAC-36 position 24,630), 2 additional Gs in a homopolymer run at BAC-36 position 25,210), and a synonymous T→C change in the K7 gene (BAC-36 position 28,409). No variant bases were found in dUMI-consensus reads at the equivalent positions, indicating that the 3 BAC-36 sequence variants were not present in this passage of the BCBL-1 line at detectable levels (i.e., <1 copy per 286 genomes).

### KSHV sequence derivation from tumor tissues and oral swabs

KSHV genome sequences were successfully obtained from samples provided by 9 participants with HIV-associated KS, including 12 KS tumors and 11 oral swabs. (**Table 1**). The representation of KSHV DNA in a sample was determined by ddPCR analysis of KSHV genes vIL-6, vBCL-2, RTA and LANA (**Fig 1A**) and provided as the percentage “on-target” KSHV DNA. These levels ranged from 0.03% to 1.35% (median 0.17%) in tumors, while most oral swab samples were below 0.01% on-target (**Table 1**). Following one enrichment with RNA baits, KSHV DNA corresponded to a median of 1.3% on-target, a >6,000-fold increase, and after a second enrichment a median of 24.2% on-target was obtained, for a total of 120,000-fold enrichment (**Table 1**).

Median total read coverage across KSHV genomes was 22,000 for tumors and 38,000 for oral swab samples. After collapsing mapped reads by dUMI, the median read coverage, corresponding to the number of viral genomes assessed, was 364 for tumors and 7 for oral swabs (**Table 1, S3A-B Fig**). Tumor sample U032-B had the highest number of genomes analyzed at 1,653. We set the lowest number of reads accepted for confident assignment of variant frequencies to be 100 (**S4A Fig**); below this number dUMI-consensus read coverage was judged to be too sparse. U020-C was an exception because its low mean dUMI-consensus read coverage was due to most of the KSHV genome being deleted, as discussed below. For other samples with mean dUMI-consensus read coverage below 80, all from oral swabs, the dUMI-consensus reads generated were insufficient to cover the entire KSHV genome, although whole KSHV genomes could be assembled from raw reads. Overall, read coverage was relatively uniform along the KSHV genome for most tumors (**S3A Fig**) and all adequately sampled oral swabs (**S3B Fig**).

Very few point mutations were found in dUMI-consensus reads from either tumors (**Fig 3A**) or oral swabs (**Fig 3B**). Excluding the major repeat regions, the number of positions with a detectable intrasample variant base ranged from 2 – 218 (<0.01 – 0.17%) (**Table 1**). These frequencies were lower or comparable to those in the BCBL-1 cell line (**Table 1**). The sample-consensus genome was generally the only KSHV sequence present in each sample, hence, there was no evidence for the existence of quasispecies [78]. However, in contrast to that observed in BCBL-1 viral genomes, clinical samples had detectable variation in long homopolymer runs.

**Figure 3.**
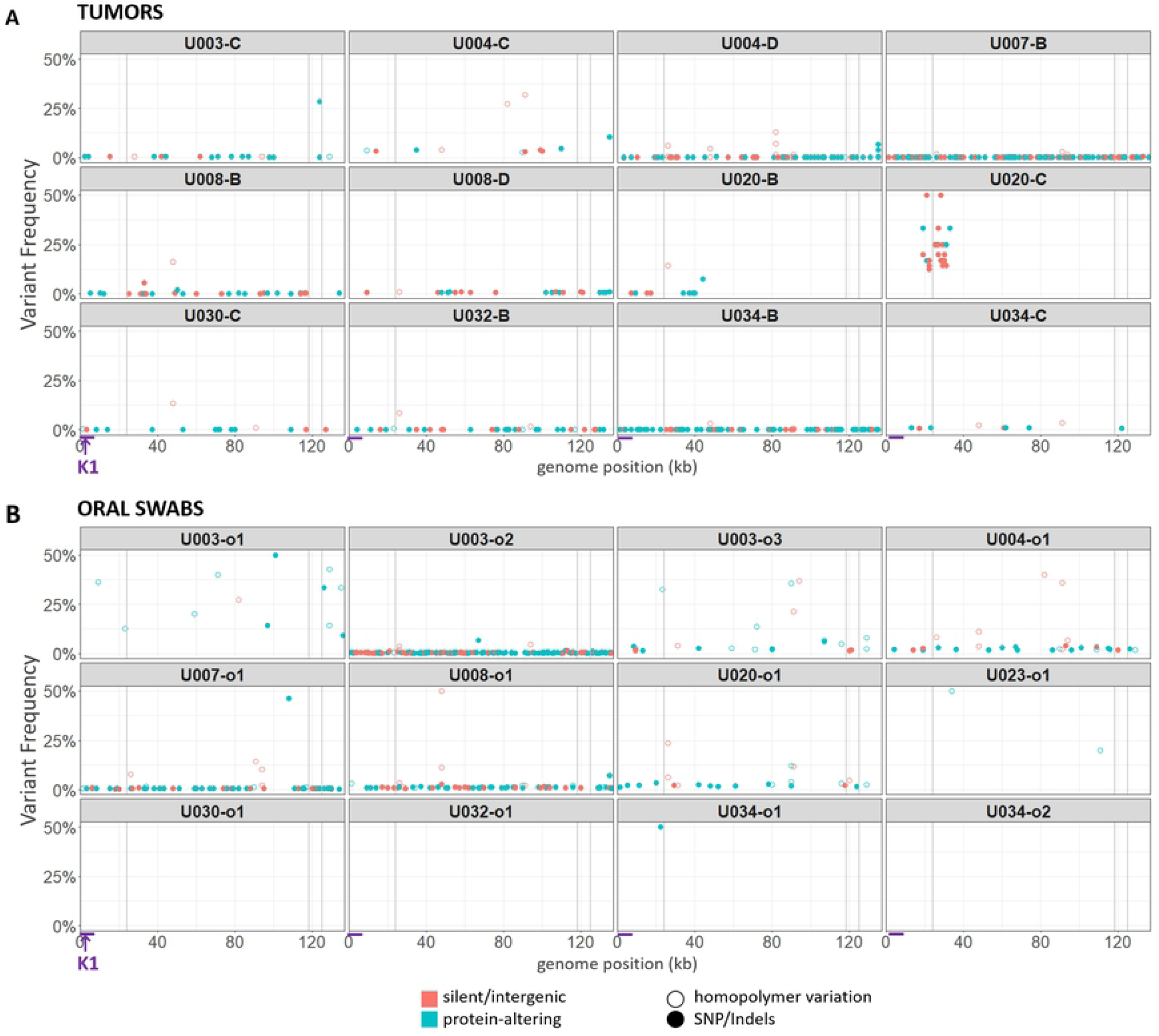
Point mutational diversity in KSHV genomes from tumors and oral swabs. Bubble plots of minor sequence variants remaining after removal of PCR errors, in KSHV genomes from tumors (A) and oral swabs (B). Each bubble represents a variant base or indel, colored by whether they were predicted to be silent or protein-altering mutations. Silent mutations include synonymous and intergenic point mutations, while protein-altering mutations included non-synonymous, nonsense and frameshift mutations. Hollow circles represent mutations occurring in homopolymer runs. Bubble heights represent the frequency of the variant base among dUMI-consensus reads at that position. Vertical gray columns represent the masked repeat regions. The region containing the K1 gene is indicated with arrows at the bottom of the figure.

Artifacts resulting from the end-repair step in DNA library preparation, which precedes the application of dUMI tags, cannot not be corrected by duplex sequencing [55,60,79]. Hence, 9 bases were removed from ends of dUMI-consensus reads before analyses, and this substantially reduced the variation observed in the raw data (data not shown). The minor base variants remaining in all samples revealed a preponderance of C→A and G→T substitutions (**S4B Fig**) as well as differences in homopolymer run lengths (**Fig 3A & B**). Most minor variants were supported by only one dUMI-consensus read. Overall, there was an inverse relationship between mean variant frequency and mean dUMI-consensus read coverage (**S4A Fig**). Thus, true minor variant frequencies could be even lower than reported here.

### KSHV genomes were virtually identical at the point mutational level between tumors and oral swab samples from the same individual

Intra-individual single nucleotide differences between tumor and oral swab samples ranged in number from 0 – 2 across the entire ∼131-kb genomes, not counting the major repeat regions. Notably, there were almost no intra-individual polymorphisms in the KSHV hypervariable gene K1 (**Fig 3A-B**). Hence, no evidence for minor KSHV variants or multi-strain infections was found in these individuals.

KSHV genomes were distinct across the 9 participants, with sequence differences ranging from 3.06-4.85%. These genomes corresponded to K1 subtypes A5, B1 and C3 (**S5A Fig**) and K15 alleles P and M (**S5B Fig**). While K1 and K15 are the most variable KSHV genes, polymorphisms along the rest of the genome have been reported to contribute more in aggregate to the total diversity of KSHV [33,34,36]. Consistent with this, maximum-likelihood phylogenetic trees using entire KSHV genomes (**S5C Fig**) were topologically distinct from those of K1 or K15. Moreover, due to numerous signatures of recombination in the evolutionary history of KSHV [36,48], differing phylogenies across sections of the KSHV genome may be better represented by a phylogenetic network (**S5D Fig)**, in which higher degrees of conflict result in a more web-like structure rather than a tree.

### Aberrant KSHV genome structures in tumors

Among the 12 tumor-derived KSHV genomes examined, 7 had anomalous read coverage that shifted abruptly once or twice along the viral genome (**S3A Fig**). In contrast, oral swab KSHV genomes from the same individuals had uniform read coverage. This argues against preferential target capture by RNA baits or other biases. Enrichment and sequencing in some were repeated, and the distinctive read coverages were reproduced. Split reads accumulated at the points of abrupt shifts in read coverage and remained after collapsing all reads by their dUMI consensus, which removes PCR chimera artifacts. Individual anomalies observed are detailed below, along with any additional evidence showing that these represented real structural aberrations in viral genomes:

### Tumor 003-C

Read coverage in U003-C was high (average of 15,635) and uniform across the KSHV genome except for a 6-bp gap within the K8.1 gene intron up to the first base of the second K8.1 exon (**Fig 4A**). No read indicated a deletion, nor was any read found with its mate pair located across the 6-bp gap. This region was PCR amplified from unsheared U003-C tumor DNA using conserved primers flanking the gap (**Fig 4B**), and no PCR product was detectable. In contrast, an intact K8.1 intron sequence was amplified and sequenced from the oral swab of the same participant (**Fig 4C**).

**Figure 4.**
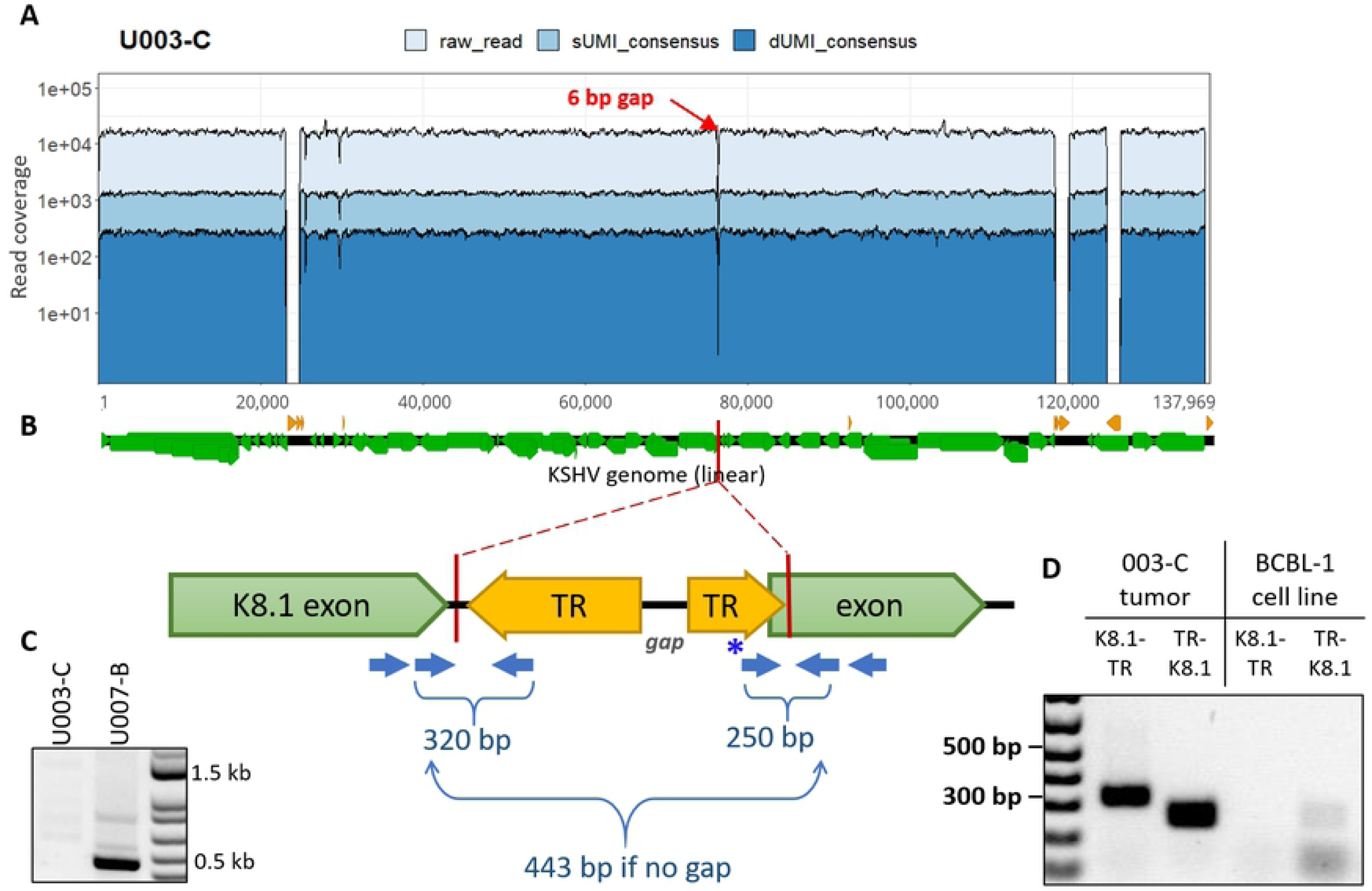
KSHV genomes in the U003-C tumor harbor a deletion within the K8.1 gene. (A) Read coverage of the U003-C KSHV genome, showing a 6-bp gap (red arrow) where no read pairs were mapped. (B) Cartoon of the *de* novo-assembly sequences generated at either side of the gap, both ended within the K8.1 gene intron and continued into terminal repeat (TR) sequences. Green and yellow arrows show the directions of the K8.1 gene and terminal repeat sequences, respectively. Blue arrows show the position of PCR primers used to confirm breakpoint junctions, with the expected PCR product sizes. (C) PCR products generated from U003-C tumor DNA using primers flanking the gap. The 443-bp PCR product expected if the K8.1 gene intron was intact was not detected from U003-C (left column), whereas the expected band was detected in tumor U007-B (right column) from another person. (D) Hemi-nested PCR of U003-C tumor DNA for the K8.1-TR (left) and TR-K8.1 (right) junctions produced products of the predicted sizes. These structures were confirmed by Sanger sequencing (data not shown). No K8.1-TR or TR-K8.1 junction fragment was produced from BCBL-1 DNA. The light bands at the TR-K8.1 lane under BCBL-1 were found to be amplicons generated from the forward primer sequence (indicated with * in panel B) overlapping with K8.1; this primer was used since the rest of the connected TR sequence was GC-rich and unsuitable for primer design.

*De novo* assembly revealed that the reverse complement of TR sequences continued from the deleted K8.1 intron sequences of U003-C (**Fig 4B**). The K8.1-TR junctions were confirmed by PCR with primers flanking the junctions (**Fig 4D**) and Sanger sequencing. Inversion of the 60-kb 3’ half of the U003-C genome, starting inside K8.1, is a parsimonious explanation for the breakpoints.

### Tumor U004-D

The first 3kb, from K1 to the end of gene ORF4, had 1.5X read coverage compared to the rest of the KSHV genome (**S3A Fig**). However, no split reads or chimeric read pairs were found to explain this result from a genome rearrangement or deletion.

### Tumor U008-B and D

U008-B had 1.7X greater read coverage over a 14.8-kb segment from inside K3 to inside ORF19 (GK18 reference positions 19,168 to 33,980, **Fig 5A**), including IR1 (masked). This was corroborated by ddPCR quantitation of vBCL-2, inside the 1.7X coverage region, with 1.7 – 1.9-fold higher gene copy number in the tumor compared to vIL-6, RTA and LANA (**Table 2**).

**Table 2.** Gene copy numbers in tumor DNA.

| Sample | vIL-6 | vBCL-2 | RTA | LANA |
| --- | --- | --- | --- | --- |
| U003-C | N/A | N/A | N/A | 9,664 |
| U003-o1 | 127 | 100 | 120 | 135 |
| U003-o2 | 1,298 | 1,238 | 1,452 | 1,628 |
| U003-o3 | 191 | 184 | 169 | 199 |
| U004-C | 1,243 | 1,274 | 1,205 | 998 |
| U004-D | 4,433 | 4,466 | 4,543 | 4,290 |
| U004-o1 | 117 | 128 | 120 | 112 |
| U007-B | 4,433 | 4,466 | 4,543 | 4,290 |
| U007-o1 | 376 | 356 | 322 | 344 |
| U008-B | 19,140 | 33,629 | 19,910 | 19,195 |
| U008-D | 24,360 | 34,755 | 12,737 | 17,189 |
| U008-o1 | 129 | 136 | 119 | 138 |
| U020-B | 49,600 | 55,850 | 4,550 | 5,500 |
| U020-C | 3,658 | 5,033 | 145 | 139 |
| U020-o1 | 254 | 231 | 234 | 248 |
| U020-o2 | 100 | 183 | 123 | 225 |
| U023-o1 | 62 | 57 | 50 | 66 |
| U030-C | N/A | N/A | N/A | 59,730 |
| U030-o1 | 2 | 5 | 5 | 7 |
| U032-B | 476 | 520 | 494 | 466 |
| U032-o1 | 9 | 0 | 2 | 0 |
| U034-B | 1,920 | 1,887 | 1,870 | 1,793 |
| U034-C | 13,083 | 13,335 | 8,148 | 6,552 |
| U034-o1 | 9 | 6 | 9 | 7 |
| U034-o2 | 6 | 11 | 0 | 1 |

N/A, did not quantify

**Figure 5.**
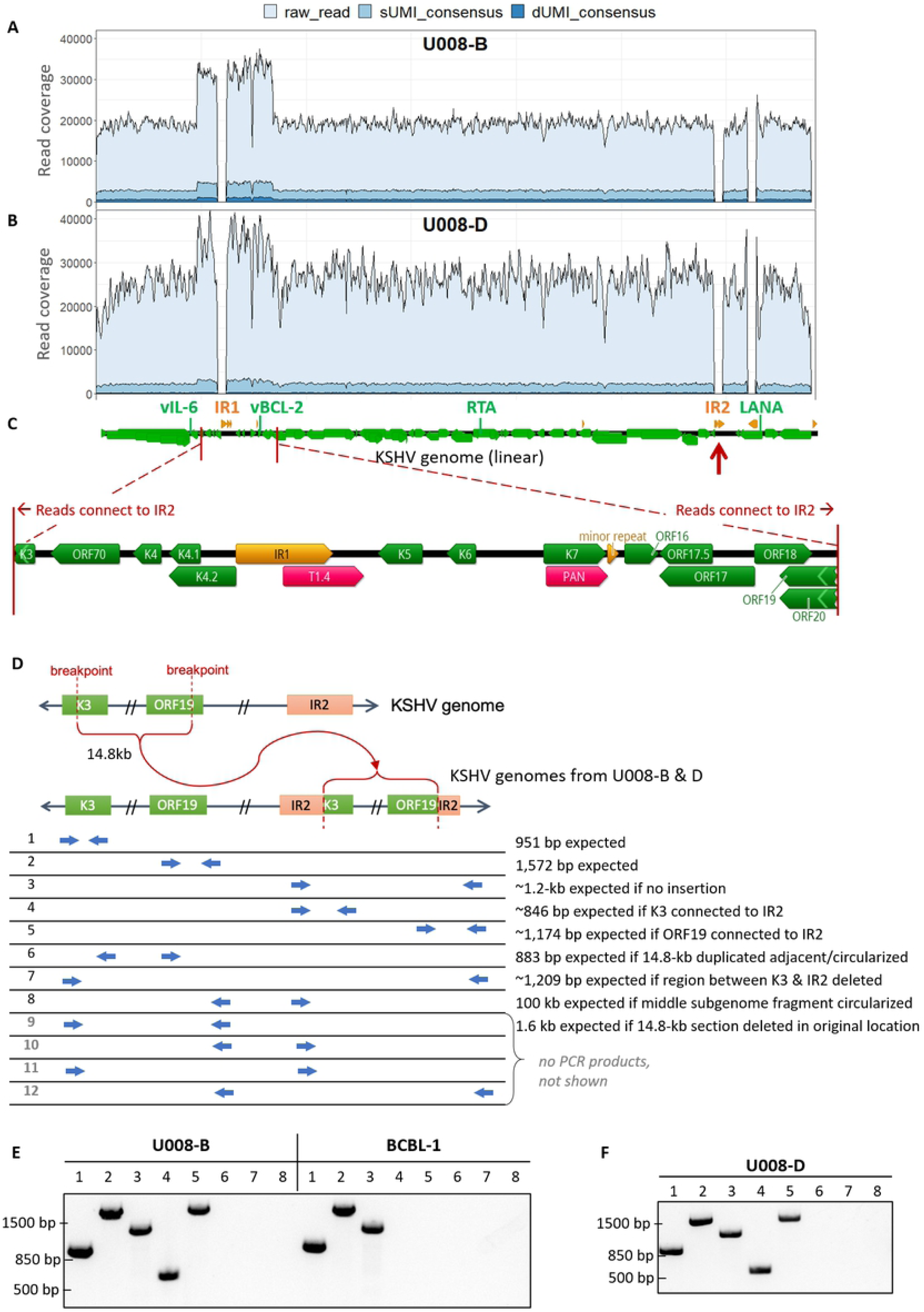
KSHV genomes in two tumors from participant U008 had a 14.8-kb region flanking Internal Repeat 1 (IR1) duplicated and translocated to Internal Repeat 2 (IR2). Total, sUMI and dUMI-consensus read coverage of tumor B (A) and D (B) genomes from individual U008. (C) Annotations of the region with 1.5-2X read coverage, with genes in green, repeat regions in orange, and long non-coding RNAs in red. Many reads on the edge of this region continue into IR2 (red arrows). Annotations are from the KSHV reference strain GK18. (D) Cartoon showing the duplication of the 14.8 kb region into IR2 and the PCR primers used to examine the genomic rearrangement in unsheared tumor DNA extracts from tumors U008-B and U008-D. PCR products produced from primer pairs numbered in D from U008-B and BCBL-1 (E) and in U008-D (F). All visible bands were excised from the agarose gel and sequenced, confirming the junction sequences. Primer pairs # 9-12 produced no PCR products discernible on an agarose gel and are not shown here.

Inferring from split reads, the 14.8-kb segment was translocated to inside IR2 (to GK18 position 119,496, **Fig 5D**). This was confirmed in the unsheared tumor DNA extract by PCR and Sanger sequencing using primers spanning the breakpoint (**Fig 5D & E**, lanes 4 & 5). Other primer pair combinations were tested to see if there were DNA species with the 14.8-kb segment inverted, deleted in place, duplicated in tandem or rearranged in other ways. None generate detectable PCR products except for primer pairs showing that the 14.8-kb segment also exists in the native configuration (**Fig 5E**). Thus, the 14.8-kb segment was copied into IR2 but had not been deleted from its original location.

In a parallel study of viral transcriptomes [80], abundant expression of a chimeric Kaposin transcript fused to the 14.8-kb segment was found in tumor U008-B, consistent with the viral genome structure we observed. Another tumor from the same participant, **U008-D** (**Fig 5B**), had 100% nucleotide identity and was confirmed to have the same duplication and breakpoint junctions (**Fig 5F**).

### Tumor U020-B

Read coverage abruptly dropped 12.8-fold over the last ∼90 kb of the KSHV genome in this tumor (**Fig 6A**). This was consistent with ddPCR quantitation, with vIL-6 and vBCL-2 gene amplicons having 9.0 – 12.3-fold higher levels than RTA and LANA (**Table 2**). The coverage shifted before the end of ORF25 (GK18 position 46,615) and reads at this breakpoint continue into TR sequences ∼90 kb downstream (**Fig 6C**). Thus, U020-B appeared to have KSHV genome variants with a ∼90-kb deletion, or formally, a 12.8X amplification of a 46-kb subgenomic region. No U020-B tumor DNA remained to allow confirmation of this breakpoint.

**Figure 6.**
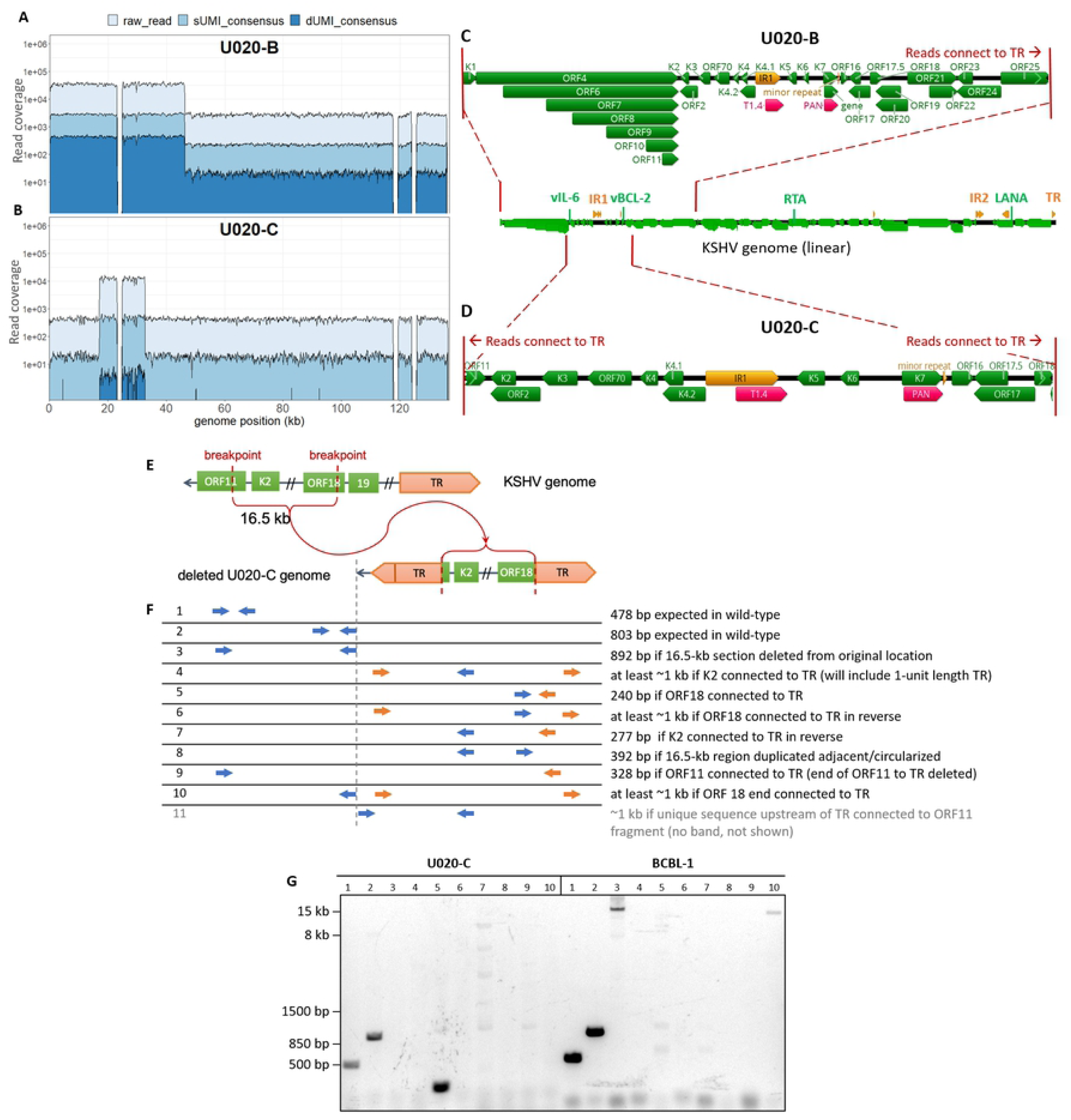
KSHV genomes in U020-B and U020-C have large, distinct deletions. Total and dUMI-consensus read coverages of U020-B (A) and U020-C (B) KSHV genomes. GK18 reference annotations in the high-coverage regions of U020-B (C) and U020-C (D). (E) Cartoon showing the region encompassing the high coverage region of U020-C viral genomes, leaving a 16.5-kb region connected to TR. (F) PCR primers used to examine the deletions. Primers to unique genomic sequences are in blue, while primers to repeat sequences are in orange. (G) PCR products produced from primer pairs numbered in E, with DNA from U020-C and BCBL-1 as templates. Bands from lanes 1, 2 and 5 were excised from the agarose gel and sequenced. Attempts to sequence the light bands in U020-C #7 and #9 were unsuccessful. Row 11 primers, in which the forward primer binds to unique genomic sequences preceding the TR, yielded no discernible product (not shown).

### Tumor U020-C

This tumor from participant U020 had (30-fold) shift in read coverage and different breakpoints inside ORF11 and ORF18 (**Fig 6B**). ddPCR quantitation demonstrated gene copy numbers of vIL-6 and vBCL-2 amplicons to be 25 – 36-fold higher than for RTA and LANA (**Table 2**). The spike in read coverage occurred over a 16.2 kb region (GK18 positions 16,942 to 33,011). Again, chimeric reads were found at either ends of this region continuing into TR sequences (**Fig 6D**), indicating fusion with the TR (**Fig 6E**). Junction fragment-specific PCR and Sanger sequencing confirmed the 3’ junction (**Fig 6G**, lane 5). No PCR product was produced from the other putative breakpoint junction, TR-ORF11 (**Fig 6F**, lane 4 primers in K2 and TR). However, the latter result is likely due to GC-rich TR sequences being largely unsuitable for primer binding. Many potential forward TR primers paired with a functional reverse TR primer (**Fig 6F**, lane 5) yielded no PCR product when a control BCBL-1 DNA was used as template.

### Tumor U030-C

A uniform >30,000 reads/position was observed throughout most of the KSHV genome. However, coverage dropped or was missing within the K15 gene (**S3A Fig**). The remaining K15 sequences corresponded to the K15 M-allele, which is less common than the P allele but was included in our RNA bait design (GenBank U75698). PCR amplification and Sanger sequencing of this region showed that the U030-C tumor did contain some copies of the entire M-allele K15 sequence. The U030-C sample-consensus genome was finished with this sequence, but no reads mapped to the gaps in K15. In the parallel RNAseq study of the same participants, transcripts of K15 were lacking in U030-B and C, despite being produced in all other tumor samples [80].

### The same aberrant KSHV genomes are found in multiple lesions from the same individual

When breakpoint junction sequences marking an aberrant KSHV genome were confirmed by PCR in a tumor, PCR primers across those breakpoints were used to screen for the same structures in other available tumors from the same individual. In the case of U008-B and U008-D, full-length genome sequencing showed that they had the same 14.8 kb subgenomic sequence duplicated in IR2 (**Fig 5F**). These two tumors were biopsied from distinct lesions on the left leg (**S6 Fig; S2 Table**). Nested PCR screening for this breakpoint junction sequence in 6 other distinct lesions (**S2 Table**) from this individual showed that no other tumors had this duplication (not shown).

In contrast, four additional tumors tested from participant U003 had the same inversion breakpoints as tumor U003-C (**Fig 7**). Moreover, no intact K8.1 sequences were detected in 2 of these 4 tumors by nested PCR of the region spanning the K8.1 intron gap (**Fig 7**). These biopsies came from distinct lesions in the left leg (**S2 Table**). Lastly, in participant U020, the ORF18-TR junction sequences found spanning the U020-C genomic deletion was not detected in the 2 other tumors tested.

**Figure 7.**
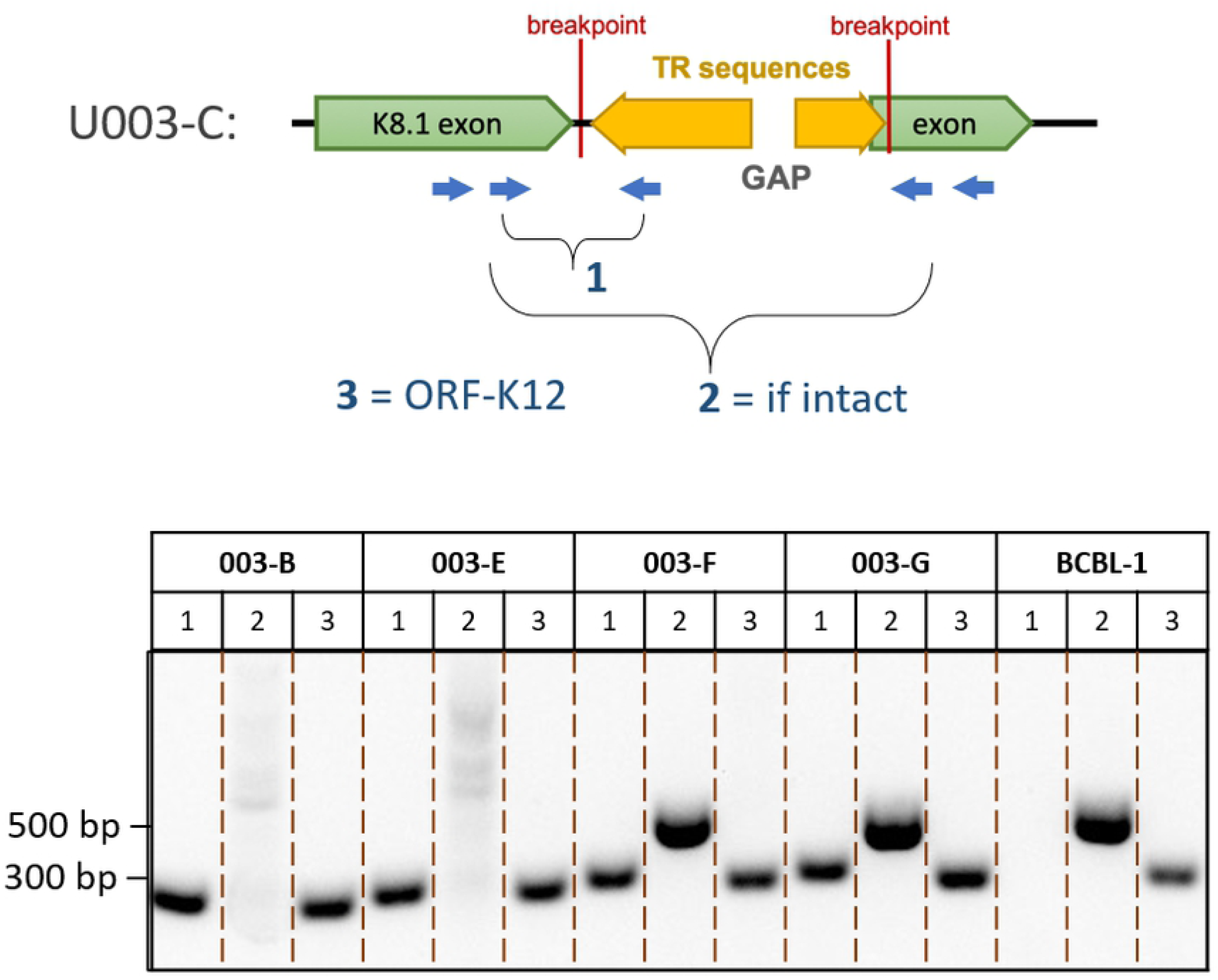
KSHV genome structures in participant U003. Junction sequences marking the genomic aberration in U003-C were detected in all 4 other tumors tested, while intact K8.1 sequences were detected in only 2. The cartoon shows the breakpoints in the K8.1 intron of U003-C extending into TR sequences, along with PCR primers used to confirm the genome structure. PCR products from other tumors of participant U003 and from the BCBL-1 cell line are shown. All visible bands were excised from the agarose gel and their structures confirmed by Sanger sequencing.

### Mutations in sample-consensus KSHV genomes from tumors impacted protein coding sequences

Among the 7 participants with KSHV sequences from at least one oral swab and one tumor, sample-consensus KSHV genomes were identical in the oral and tumor samples of 2 participants and differed in 4 others. In the remaining participant, U004, the sample-consensus KSHV genome in one tumor was identical to that in oral but the second tumor had mutations. The mutations unique to tumors were typically nonsynonymous point mutations resulting in highly dissimilar residues or other mutations likely to disrupt their expression (**Table 3**).

**Table 3.** Unique KSHV mutations observed in tumors compared to oral swabs from the same individual

| Sample ID | Tumor-specific differences |
| --- | --- |
| **U003-C** | K12 synonymous mutation within miR-K10<br>genomic inversion starting at K8.1 |
| **U004-C**<br>**U004-D** | NONE<br>ORF32 nonsynonymous mutation R56Q<br>K15 nonsynonymous mutation A290P<br>28-nt deletion within the K8.1 promoter<br>3-kb segment duplication from before K1 to after ORF4 |
| **U007-B** | NONE |
| **U008-B**<br>**U008-D** | duplication of 14.8 kb segment around IR1 into IR2<br>- breakpoints inside K3 & ORF19<br>- genes duplicated: ORF70, K4.1, K4.2, K5, K6, K7, ORF16, ORF17, ORF17.5, ORF18<br>same as U008-B |
| **U020-B**<br>**U020-C** | ORF25 nonsynonymous mutation Q594K<br>genomic deletion connecting end of ORF25 coding sequence to TR sequences, 47 kb remaining<br>ORF11 nonsynonymous mutation T396P<br>K3 nonsynonymous mutation F88L in transmembrane domain<br>K8.1 nonsense mutation at start of 2nd exon<br>genomic deletion leaving only 16 kb segment surrounding IR1 connected to TR sequences<br>- breakpoints inside ORF11 and ORF18<br>- ~30X coverage for: K2, ORF2, K3, ORF70, K4, K4.1, K4.2, K5, K6, K7, ORF16, ORF7, ORF17.5 |
| **U032-B** | ORF63 nonsynonymous mutation T848A |
| **U034-B**<br>**U034-C** | NONE<br>NONE |

Several tumor-unique mutations or genome aberrations occurred in structural genes (**S3 Table**), and frequently involved the K8.1 gene, which encodes an envelope glycoprotein. The U003 inversion breakpoint cleaved the K8.1 gene. U004-D had an R56Q mutation in its ORF32 tegument protein coding sequence, as well as a 28-nt deletion in the promoter region of K8.1 (**S7A Fig**). The deletion was after the K8.1 core promoter sequence [81], but encompassed the K8.1 transcription start site [82]. The ORF25 major capsid protein in U020-B had a Q594K mutation, in addition to the U020-B genomic deletion that started downstream of ORF25. U020-C had a nonsense mutation at the beginning of the second K8.1 exon. Finally, U032-B had a T848A mutation in ORF63, a tegument protein.

The only intra-host synonymous point mutation observed was in ORF K12 of U003-C (GK18 position 118,082). This C to T change occurred within the oncogenic microRNA K10 (miR-K10) sequence in the Kaposin A transcript. The three oral swab samples from this participant maintained the consensus C at this position (**S7B Fig**), whereas the 4 other tumors from this participant examined had T at this position, with tumor U003-G having a minor population of viruses with the consensus C (**S7C Fig**). Among published KSHV genomes, only ZM106 (GenBank KT271458), also derived from a KS tumor, had a T at this position.

### Lack of evidence for integration of KSHV sequences into human chromosomes

No *de novo*-assembled scaffolds, split reads or improperly-paired read mappings suggested any instance of KSHV sequences fused to human DNA. Nevertheless, attempts were made to systematically search for human-KSHV chimeric sequences. The methods employed were those used to screen for all integrated herpesviruses sequences in public databases [83] and EBV integration sites in in primary gastric and nasopharyngeal carcinomas [84]. The KSHV genome inversions, duplications and deletions described above were detected by LUMPY with high confidence values. In contrast, putative breakpoints that joined human and KSHV sequences were supported by only tens of reads, about 2 orders of magnitude lower in number, and often involved LANA repeats into low-complexity human repeat sequences.

### Co-infection with EBV detected predominantly in oral swabs

Some scaffolds during *de novo* assembly correspond to EBV sequences. Nearly all oral swab samples yielded multiple EBV-mapping scaffolds up to 73 kb, with no region of the EBV genome over-represented. In contrast, EBV-sequences were detected in only 5 of 12 tumors, and in all cases were sequences flanking the EBNA-1 repeat (**S8 Fig**). The proportion of reads mapping to EBV in oral swabs ranged from 0 - 33%, median 1.8%, whereas in tumors the range was from 0 - 0.5%, median 0.002% (**S4 Table**). No other eukaryote viruses were identified, including HIV, with which every participant was known to be infected.

## Discussion

This study is the first to explore KSHV intra-host diversity at the whole genome level and provided an unprecedented level of precision to herpesvirus genome sequence analysis in clinical specimens, with >100 essentially error-free genome sequences obtained from most tumors. KSHV genomes were obtained from 11 oral swabs and 12 KS tumors of 9 Ugandan adults with KS. By incorporating dUMI, PCR misincorporation errors and template switching artifacts were substantially eliminated, permitting detection variants as infrequent as 0.01% and a theoretical error rate of 1/10^−9^, or approximately the DNA replication error rate in eukaryotic cells [55].

There were no signs of KSHV quasispecies, consistent with large dsDNA viruses having the lowest mutation rates among viruses [85]. Less than 0.01% of all base positions in the 131-kb KSHV genomes (excluding the major repeat regions) were found to have a detectable variant, typically supported by only one dUMI-consensus read. This exceedingly low intra-sample variation is within the published resolution of duplex sequencing [55]. While there are reports of intra-host KSHV variability in certain KSHV-endemic populations [38], in children [43], in iatrogenic settings [39–41] and in blood of AIDS-KS patients [42], these findings were arrived at by Sanger sequencing of PCR amplicon clones of hypervariable regions in K1 or other genes. Such protocols are more likely to detect errors that occurred during PCR. Our study found virtually no intra-sample or intra-host diversity even at K1 in the 9 individuals examined. Recombination is evident in the evolutionary history of KSHV [36,44,48,49]; hence, co-infections by multiple KSHV strains must occur, if sporadically.

The most striking observation in this study was the frequency and tumor-specificity of aberrant KSHV genomes, summarized in **Figure 8**. Up to 7 of the 12 KS tumors examined had major inversions, deletions or duplications comprising the majority of KSHV genomes in those tumors. In stark contrast, no aberrant genome structures were found in oral swabs. It is unclear whether the tumor-specific mutations observed were required for tumorigenesis or tumor persistence, or whether they were a consequence of localized genomic instability known to occur in tumors [86], but several observations suggest that these changes were not random. Rearrangement breakpoints and other mutations almost always occurred inside coding sequences of lytic genes (**Tables 3 & S3**), with many truncating protein coding sequences. These mutations may have been selected for, if for instance expressing these proteins exposed host cells to immune targeting. Mutations that disrupt genes may also indicate that those genes were not necessary for sustaining tumorigenic growth. Conversely, regions of the KSHV genome that were duplicated, conspicuously intact or translocated next to strong promoters (as in 008-B and 008-D) may point to KSHV genes that are influential in driving tumor cell proliferation.

**Figure 8.**
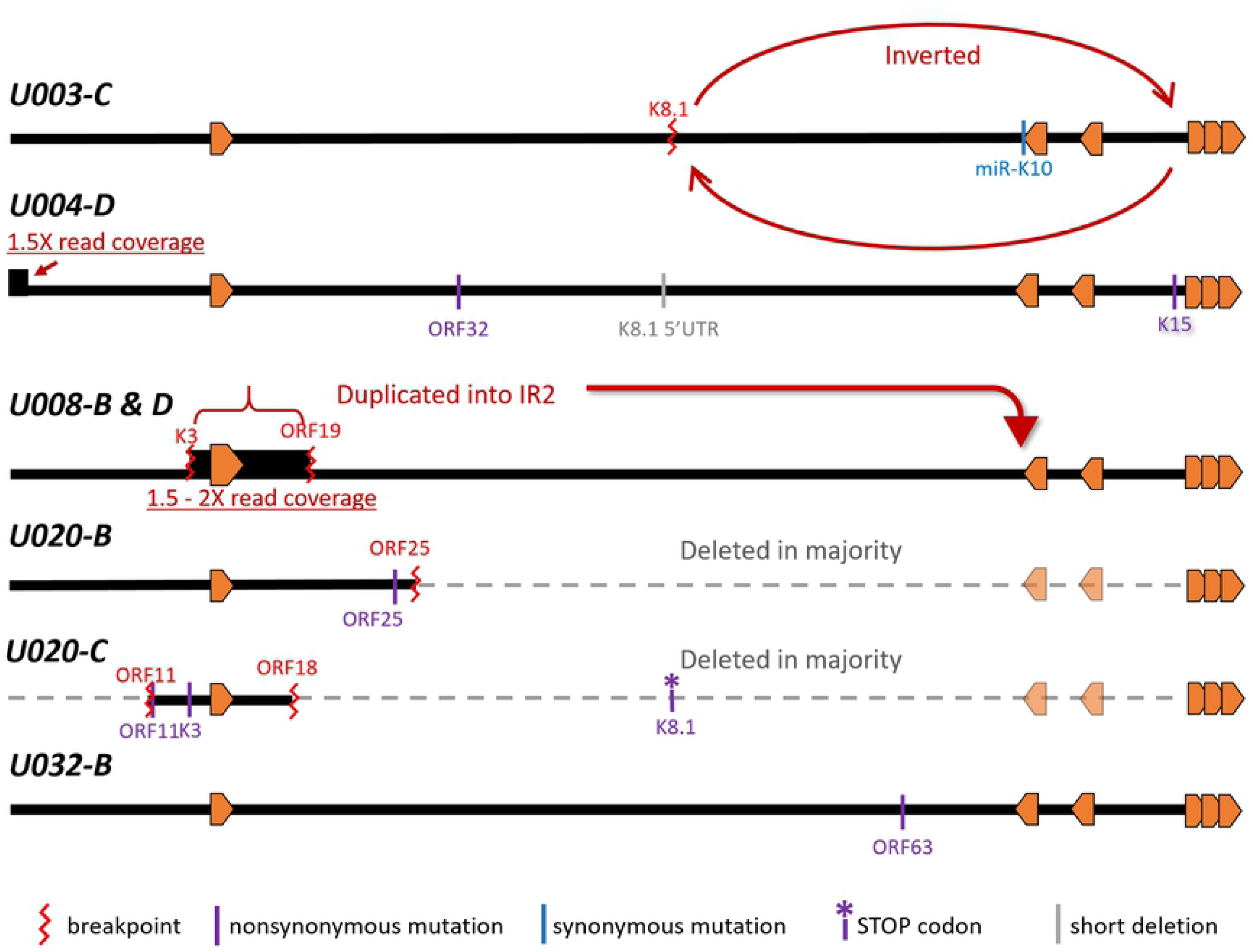
Schematic representation of the 5 aberrant KSHV genomes discovered in KS tumors. Specific details and evidence for each are referred to in the text and in Table 3.

The genomic region around IR1 featured prominently in genomic rearrangements in 4 tumors, potentially leading to their over-expression relative to other KSHV genes. For example, tumors U008-B and U008-D had a 14.8-kb portion of their genomes, from inside K3 to ORF19, duplicated into within IR2 (**Fig 5**). In a parallel RNAseq study, tumor U008-B had been found to abundantly express a chimeric transcript of the 14.8 kb section fused to IR2 sequences transcribed from a strong latency-associated promoter [80]. Distinct deletions were observed in tumors U020-B and U020-C from another participant, but the genomic regions retained, aside from TR sequences, again included the IR1 region (**Fig 6**). There is evidence to suggest that KSHV lytic gene expression is crucial to KS pathogenesis [87], and that residual lytic gene expression plays a role in latent KSHV persistence *in vivo* [88]. IR1 is one of the origins of lytic replication, and transcripts around IR1 are among the most highly expressed in KS tumors [80]. These include two long non-coding RNAs that have indispensable roles during lytic reactivation of KSHV, T1.4 [67,89,90] and PAN [91–93]. PAN has been shown to interact with promoters of cellular genes involved in inflammation, cell cycle regulation and metabolism, and exogenous expression of PAN alone enhanced cell growth phenotypes [94]. Recently, virally-encoded circular RNAs encoded within PAN were discovered to be abundant in clinical samples and were inducible in KSHV-infected cell lines [95–97]. Other non-coding transcripts are potentially expressed from this region but their biological significance is unknown [93,98]. Finally, most ORFs encoded in the 14.8-kb retained region are intermediate-early or delayed-early lytic genes that may have functions in subverting adaptive (K5/MIR2) [99,100] or innate immunity (K4/vCCL-2, K4.1/vCCL-3, K4.2 and K6/vCCL-1)[100–103], and apoptosis (K7 and ORF16/vBCL-2) [100,103].

Other than genomic rearrangements, sample-consensus KSHV genomes in tumors and oral swabs within the same individuals differed by at most two point mutations or a short deletion. No mutations occurred in intergenic regions, and almost all were nonsynonymous changes resulting in highly dissimilar amino acids. Notably, the sole intra-host synonymous mutation found occurred inside K12/Kaposin A of tumor U003-C (GK18 position 118,082), within the embedded microRNA miR-K10. The oral swab counterpart maintained the database consensus. Expression of the Kaposin A transcript is tumorigenic [104] and a single base change has been observed to abolish this effect [105], although a different mutation was observed here.

The late lytic gene K8.1 was found to be mutated in KS tumors from 3 individuals. U003-C had an inversion breakpoint at the K8.1 intron, U004-D had a 28-bp deletion ending at 4 bases upstream of the first K8.1 exon, and U020-C had a nonsense mutation at the start of the second K8.1 exon (**Fig 8**). Furthermore, intact K8.1 gene sequences were undetectable by PCR in most of participant 003’s tumors tested (**Figs 4C & 7**). Truncations in K8.1 had been reported previously, all from KS tumor isolates. The original GK18 isolate had a 74-bp deletion at the 3’ end (see GenBank ID AF148805 K8.1 annotation); the Zambian isolate ZM124 (GenBank ID: KT271466) had a 25-nt deletion resulting in a frameshift and premature stop [33]; finally, Japanese isolate Miyako1 has a stop codon early in its first exon (see GenBank ID LC200586 miscellaneous annotation). Collectively, these findings suggest selection pressure against K8.1 expression in tumors.

Gene K8.1 encodes an envelope glycoprotein that interacts with heparin sulfate for attachment [106–109]. It is not required for entry into endothelial [107] or 293 cells [110], although it had recently been shown to be necessary for infection of primary and cultured B-cells [111]. The K8.1 protein is often used as an indicator of the late lytic stage and is among the most immunogenic KSHV proteins [112–114]. It is therefore conceivable that the preponderance of K8.1 mutations might be due to potent immune targeting of cells expressing K8.1 glycoproteins.

Some of the aberrant KSHV genomes we observed would be unable to produce infectious virions, yet the same viral genome rearrangements were sometimes found in multiple lesions. All 5 tumors tested of Participant U003 had the K8.1-TR junction present, but intact K8.1 sequence was detectable by PCR in only 3 (**Fig 7**). Participant U008 had the same sequence duplication in 2 distinct tumor lesions (**Figs 5 & S6)**. Thus, spread of these mutated genomes could have occurred by tumor metastasis or with a helper virus.

Detection of aberrant KSHV genomes is not without precedent. The first whole genome sequence of KSHV published reported a 33-kb portion of the KSHV unique central region duplicated into the TR region [115]. A study of 16 tumor-derived KSHV whole genomes from Zambia reported 4 that had regions with 3-fold more coverage than the sample average, although these were not examined [33]. A PCR screen for some KSHV genes showed that some KS tumors and KSHV-infected B-cell lines can harbor deleted KSHV genomes [116], and one such B-cell line proliferated faster than the parental BCBL-1 line [116]. The infecting KSHV had an 82-kb deletion from the 5’ end of its genome, was lytic replication-incompetent, and could be packaged by a helper virus.

The LANA protein tethers the KSHV episome and is required for maintaining KSHV latency and for latent replication of TR sequence-containing plasmids [117,118], but the deleted 020-B and 020-C genomes had the entire latency locus, including LANA, missing. Remaining intact KSHV genomes in the same cell could be supplying LANA, as latently infected cells frequently harbor multiple KSHV copies per cell [118,119], and KSHV episomes are inherited by daughter cells in clusters [120].

Two remaining observations of note in this study were that, in contrast to larger studies of EBV in tumors [84,121], we found no integrations of KSHV into human chromosomal DNA. Secondly, our cohort was infected also with EBV, as seen in the abundance of sequence reads corresponding to EBV especially in oral swabs. Tellingly, the rare EBV sequences found in tumors were consistently of EBNA-1, whose GC-rich repeat domain has nucleotide homology to LANA repeats [122] and hence were probably co-captured by the RNA baits. No sequences of HIV or other eukaryote viruses were detected.

In summary, highly accurate deep sequencing of whole KSHV genomes in paired oral swab and KS tumors from individuals with advanced KS were virtually identical at the point mutational level. Where there were differences, the oral viruses had the database consensus genotype while tumor viruses had novel mutations. KS tumors can harbor KSHV with genomic aberrations or other mutations that may alter lytic gene expression, and these viral mutations can be shared by distinct KS tumors within an individual. Our study points to associations with KS tumors of the region surrounding IR1 and the K8.1 gene. As inactivating mutations seem to be a frequent feature of tumor-derived KSHV, whole genome or targeted sequencing may reveal more viral genomic regions important to the pathogenesis or persistence of KS.

## Acknowledgements

We thank the study participants who contributed invaluable specimens that make this study possible. We also thank D. Depledge and J. Breuer for sharing their RNA bait sequences for capturing KSHV genomic sequences and M. Lagunoff for kindly providing us with an early passage BCBL-1 cell line.

## Figure Legends

**S1 Figure. dUMI-adaptors and primers for duplex sequencing.** During library preparation, sheared DNA fragments were A-tailed and ligated with forked, double-stranded oligonucleotides containing Illumina TruSeq universal adaptor sequences, 12-random base pairs as dUMI and spacer sequences. The adapted DNA libraries were PCR amplified before enrichment with primers mws13 and mws20, which bind to Illumina Truseq adaptors. Primer mws21 containing sample index ID for multiplex sequencing was used for PCR following enrichment. DNA libraries post processing are shown at the bottom.

**S2 Figure. Workflow of genome assembly and variant analysis.** KSHV genomes were first assembled *de novo* from sequence reads of each sample, before being used as reference for mapping their respective dUMI-consensus reads. See details in the Methods section. Discrepancies in bases between the sample-consensus genome and mapped dUMI-consensus reads were taken to be real intra-sample variants.

**S3 Figure. Read coverage.** Raw (light blue), sUMI (blue) and dUMI-consensus (dark blue) read coverage in log scale along the de novo assembled, sample-consensus KSHV genomes in tumors (A) and oral swabs (B) examined in this study. Major repeat regions were masked with inserted Ns in all sample-consensus genomes.

**S4 Figure. Potential sequencing artifacts.** (A) KSHV intrasample variant frequency as a function of read coverage. Sample variant frequencies were estimated (Table 1) when at least 100 viral genomes were sampled. (B) Minor variants detected in dUMI-consensus reads of all samples, by type of base substitution.

**S5 Figure. KSHV phylogenetic relationships by variable regions K1 and K15 and by whole genomes.** Phylogenetic trees of (A) K1 genes, (B) K15 genes and (C) whole genomes from this study and of select genomes from other publications. K1 and K15 subtypes are indicated to the right of the K1 (A) and K15 (B) trees. (D) A neighbor-net phylogenetic network of all published KSHV genomes to date, color-coded by genome types proposed in [36]: P1 in green, P2 in blue, N in purple, M1 in red and M2 in maroon. All *de novo*-assembled genomes from this study are in bolded italics.

**S6 Figure. U008-B and U008-D were from distinct lesions on the left leg.** U008-B biopsy was obtained from lesions in the upper thigh, while U008-D was biopsied from a large lesion on the knee.

**S7 Figure. Mutations of KSHV genomes in tumors from participants 004 and 003.** (A) Alignment of KSHV genomes from Participant 004, showing a 28-bp deletion in the K8.1 promoter in U004-D. U004-D and U004-C are from tumors while U004-o1 is from an oral swab. (B) The only intra-host synonymous mutation found in this study, within miR-K10 in participant 003. (C) Sequence chromatograms of miR-K10 in other tumors of participant U003, with a T in all tumors and a mixture of T and the database consensus C in a minority of viruses in U003-G.

**S8 Figure. Scaffolds of EBV sequences produced in 4 KS tumors map to the EBNA-1 gene.** A portion of the EBV genome is illustrated with an example of the 2 scaffolds (grey striped bars) generated from the reads of 4 KS tumors. Vertical stripes inside the gray bars indicate mismatches to the EBV reference (GenBank ID: DQ279927).

